# RF heating of deep brain stimulation implants in open-bore vertical MRI systems

**DOI:** 10.1101/650960

**Authors:** Laleh Golestanirad, David Lampman, Ehsan Kazemivalipour, Hideta Habara, Ergin Atalar, Joshua Rosenow, Julie Pilitsis, John Kirsch

## Abstract

**Purpose:** Patients with deep brain stimulation (DBS) implants highly benefit from MRI, however access to MRI is restricted for these patients due to safety hazards associated with RF heating of the implant. To date, all MRI studies on RF heating of medical implants have been performed in horizontal closed-bore systems. Vertical MRI scanners have a fundamentally different distribution of electric and magnetic fields and are now available at 1.2T, capable of high-resolution structural and functional MRI. This work presents the first simulation study of RF heating of DBS implants in high-field vertical scanners.

**Methods:** We performed finite element electromagnetic simulations to calculate SAR at tips of DBS leads during MRI in a commercially available 1.2 T vertical coil compared to a 1.5 T horizontal scanner. Both isolated leads and fully implanted systems were included.

**Results:** We found 10-30-fold reduction in SAR implication at tips of isolated DBS leads, and up to 19-fold SAR reduction at tips of leads in fully implanted systems in vertical coils compared to horizontal birdcage coils.

**Conclusions:** If confirmed in larger patient cohorts and verified experimentally, this result can open the door to plethora of structural and functional MRI applications to guide, interpret, and advance DBS therapy.

## Introduction

Deep brain stimulation (DBS) is a neurosurgical procedure that uses an implantable pulse generator (IPG) in the chest to send electric pulses via subcutaneous leads to specific nuclei in the brain to modulate their behavior. DBS has FDA approval for treatment of Parkinson’s disease, essential tremor, and epilepsy [1, 2], and a humanitarian device exemption for treatment of dystonia and obsessive compulsive disorder [3]. New studies show efficacy of DBS for an expanding range of neurologic and psychiatric disorders including Tourette’s Syndrome [4, 5], bipolar disorder [6], Schizophrenia [7], and depression [8, 9]. The rapid increase in indications of use and application of DBS parallels the large availability and need for MRI. There is a consensus that meticulous application of neuroimaging, both for target verification and for postoperative monitoring of stimulation effects is essential to rule out complications, interpret clinical outcomes, and design enhanced therapeutic protocols. When investigating the neuromodulatory effects of DBS, neuroimaging studies have largely used positron emission tomography (PET) or single photon emission tomography [10–15]. MRI has clear advantages over both PET and single photon emission tomography due to its excellent soft-tissue contrast, non-invasive nature, and the richness of post-processing analytical methods that are available to use. The clinical community however, has been cautious in adopting MRI for DBS patients due to safety concerns associated with radiofrequency (RF) heating of implants. The major safety hazard is the “antenna effect”, where the electric field of MRI transmit coil induces RF currents on implanted wires, leading to an increase in the specific absorption rate (SAR) of energy deposition in the tissue. Serious injuries have underscored the severity of such RF heating [16]. Consequently, the conditions under which DBS patients are indicated for MRI are limited. For example, under current guidelines either the whole-head SAR of the pulse sequence should be less than 0.1 W/kg (30 times below the FDA limit for imaging in the absence of implant) or the rms of B_1_^+^ field should be less than 2µT, a limit well surpassed in most clinical applications [17, 18]. Current MR labeling of DBS devices however, is limited to horizontal (closed-bore) MRI systems. Vertical MRI scanners originally introduced as “open” low-field MRI systems are now available at 1.2T. To date, no DBS SAR literature exists for this class of scanners, which are now available at field-strengths capable of high-resolution structural and functional studies. Vertical MRI scanners have 90° rotated coils which generate a fundamentally different electric and magnetic field distribution. It is well established that the orientation and phase of MRI incident electric field along the trajectory of elongated implants plays an important role in determining the magnitude of SAR amplification around exposed tips of the implant [19–24]. Therefore, it is possible that this class of MRI scanners offers a whole new solution to the problem of safe DBS imaging which has been overlooked.

This work presents the first simulation study of RF heating of DBS implants in vertical MRI scanners. Patient-derived models of DBS systems representing three major clinical device configurations were constructed from patient imaging data and simulated in a commercial open-bore 1.2T MRI system (OASIS, Hitachi) and a horizontal 1.5T body birdcage. We found a 20-30-fold reduction in SAR amplification at tips of DBS leads during RF exposure in vertical coil compared to horizontal birdcage in both isolated and fully-implanted DBS systems.

## Methods

### Patient models

DBS surgery is typically performed in two stages. In the first stage leads are implanted in targeted brain nuclei and the other end of the lead is tucked under the scalp for later connection to the neurostimulator (Figure 1, Patient 1). MRI at this stage is useful to rule out complications and to verify the target. In the next stage, an IPG is implanted in the chest and leads are connected to the IPG via subcutaneous extensions (Figure 1, Patients 2&3). Patients may receive two single-channel IPGs implanted bilaterally in the pectoral region, each stimulating one lead (Figure 1, Patient 2), or one double-channel IPG to stimulate both left and right leads (Figure 1, Patient 3). The majority of DBS patients who need MRI for clinical assessment have fully-implanted devices. MRI of fully-implanted DBS systems is also highly desired to perform functional studies that investigate changes in patterns of brain activation as a function of stimulation parameters [25, 26].

**Figure 1:**
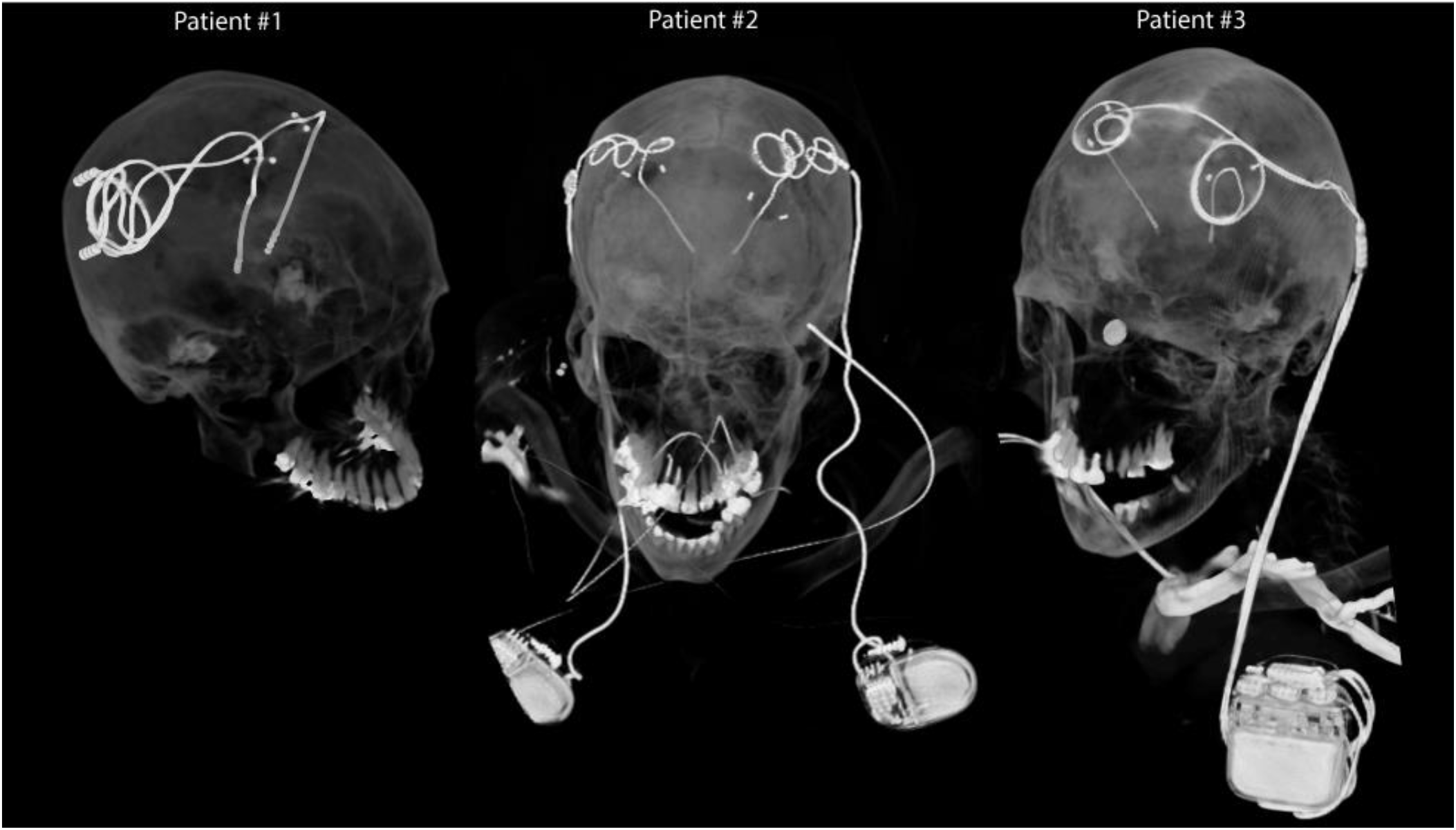
Postoperative CT images of patients with isolated and fully-implanted DBS systems used in simulations

To account for major clinically-relevant DBS configurations, we simulated one representative patient of each category. Secondary use of imaging data for simulation and modeling was approved by the ethics review board of Northwestern University and by the IRB at Albany Medical College. Postoperative CT images of three patients who had gone through subthalamic nucleus (STN) DBS surgery were used to reconstruct 3D trajectories of the implants. Lead trajectories were extracted from post-operative CT images and 3D models of leads, extensions and pulse generators were constructed following a procedure described in our previous works [23, 27, 28]. Briefly, an image segmentation tool (Amira, Thermo Fisher Scientific, Waltham MA) was used to extract the hyper-dense trajectory of DBS implants using a thresholding algorithm and preliminary 3D surfaces of the leads, extensions and IPGs were constructed. The 3D surfaces were exported to a CAD tool (Rhino3D®, Robert McNeal and Associates, Seattle, WA) in which lead trajectory lines were manually reconstructed and models of electrode contacts, core, insulation, and the IPG were created around them. Figure 2 shows patient models and implant details as used in finite element simulations.

**Figure 2:**
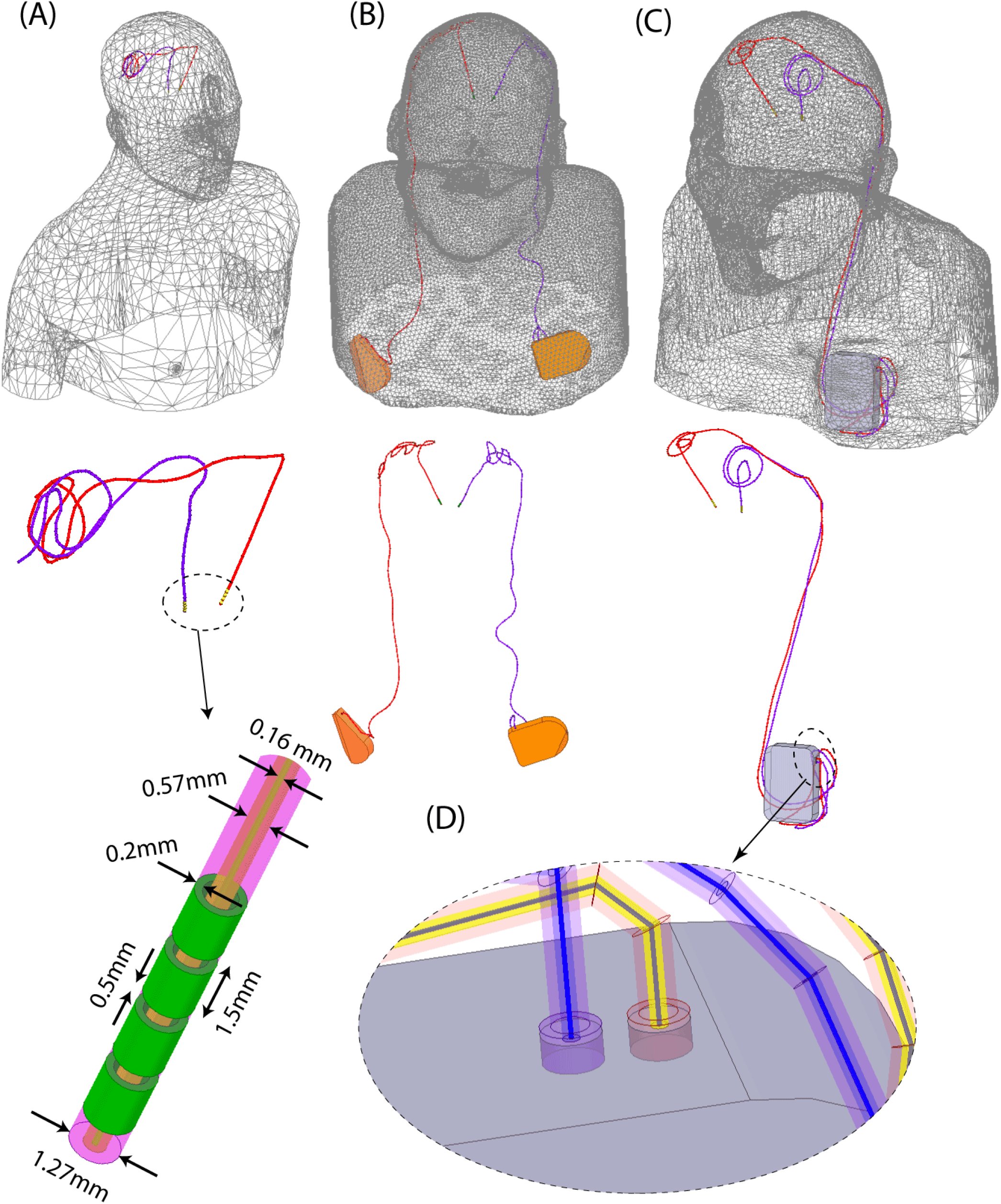
Patient models developed for finite element simulations.

Leads were composed of four cylindrical electrode contacts connected through a solid straight core made of platinum-iridium (Pt:Ir *σ* = 4 × 10^6^S/m), encapsulated in a urethane insulation (D=1.27mm, *ε*_*r*_ = 3.5) with an air-filled lumen (D=570µm). The IPG was modelled as a solid box of platinum-iridium. There was a 0.5mm gap between the end of the metal core of the extension cable and the conductive face of the IPG, representing an open circuit. Homogeneous body models (*ε*_*r*_ = 80, *σ* = 0.47 S/m) were constructed from silhouettes of patients 2 and 3 based on CT images. For patient 1, who did not have head and neck CT available, we registered DBS lead models to a standard homogeneous body model of head and upper chest.

### RF coils and field calculations

Models of a 12-rung radial flat vertical birdcage coil tuned to 50.35 MHz (1.2T) and a generic solenoidal horizontal birdcage tuned to 64 MHz (1.5T) were constructed and used in simulations. The vertical coil was designed based on a commercial coil described in [29]. Figure 3 gives geometrical details of the coil models, as well as plots of their magnetic field on a central coronal plane passing through the body of patient 2 in the absence of the implant. The counter clockwise rotating component of the magnetic field vector was calculated as 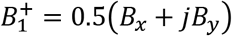. For all simulations, the input power of coils was adjusted to generate an average 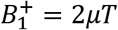 on an axial circular plane (radius=5cm) positioned 20 mm below the distal tip of DBS implants. The location of this plane was chosen to allow an unbiased averaging of 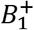 for both coils free of implant-induced distortion. 1g-averaged SAR (SAR1g) was calculated using HFSS built-in SAR module that implements IEEE STD P1528.4 recommendation [30]. The maximum of SAR1g was recorded inside a cubic area of 20mm× 20mm× 20mm surrounding four electrode contacts of the lead and used to compare two coils. Simulations were performed using ANSYS Electronic Desktop (HFSS 19.2.0, Designer, ANSYS Inc., Canonsburg, PA).

**Figure 3:**
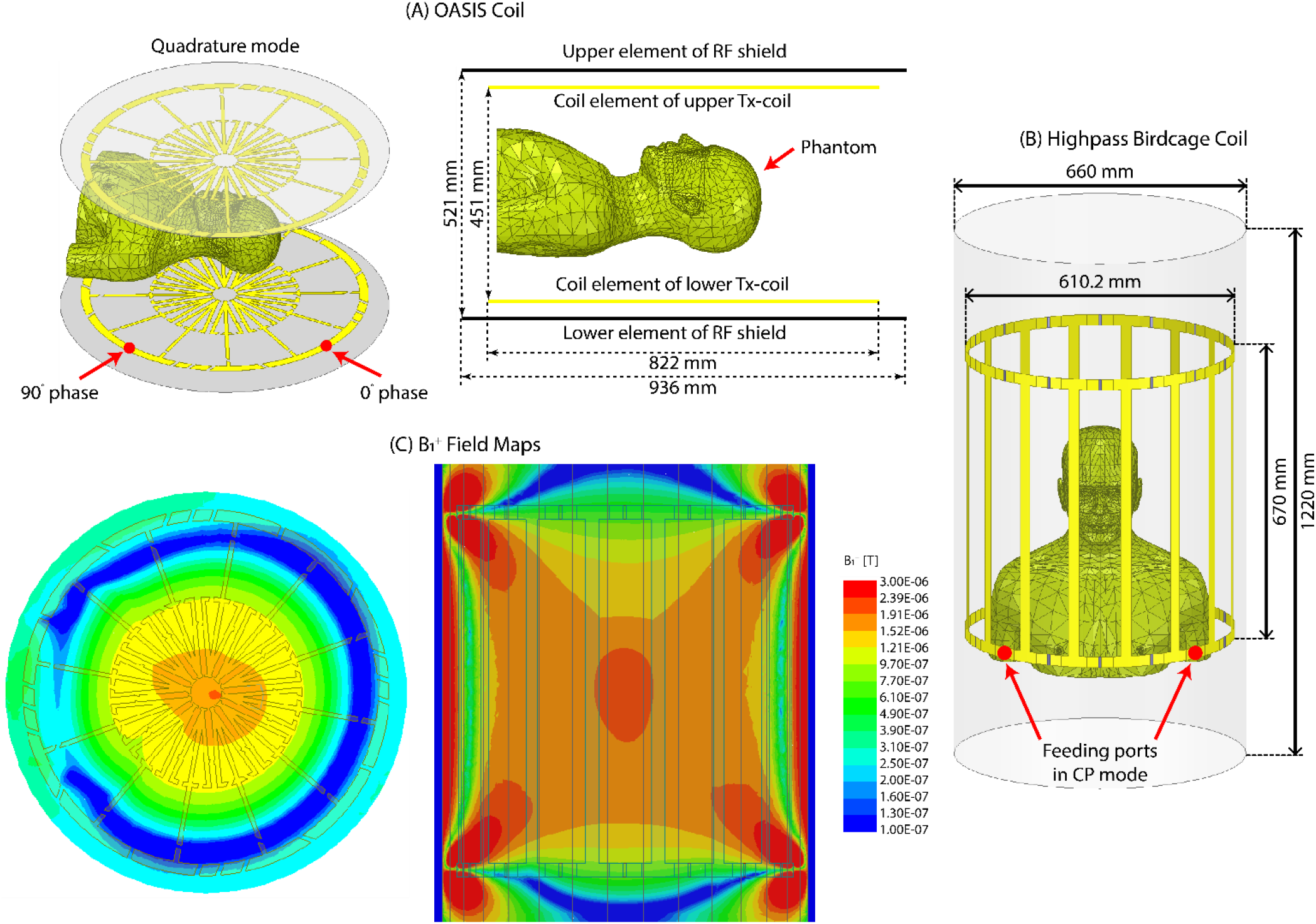
Geometry of (A) a butterfly vertical coil based on a commercially available prototype (OASIS, Hitachi) [29] and (B) a generic high-pass birdcage coil with dimensions reported in the literature [31]. (C) B1+ field maps on a coronal plane.

### Incident electric field

Recently we demonstrated that the magnitude of the tangential component of MRI electric field along the first few centimeters of extracranial trajectory of DBS leads can be used as a predictor of severity of SAR amplification at electrode tips [23]. This observation was not surprising, considering incident electric field is responsible to induce RF currents on elongated conductive leads which eventually dissipate in the tissue and cause heating. To compare the orientation of the incident electric field of horizontal birdcage coils vs the vertical coil with respect to patients’ lead trajectories, we calculated the incident E_tan_ along the length of leads and extensions using the method described in our previous work [23]. In brief, we reran simulations without implant being present but with polylines representing leads and extension trajectories imported to HFSS. We then extracted the unit tangent vectors along these lines (â) and calculated E_tan_(t) at each point along the length of the lead as:

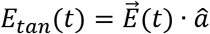

where 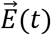 is the incident electric field of the coil and â is the unit vector tangential to the lead’s trajectory.

## Results

Figure 4 shows the distribution of local SAR around DBS lead tips of patients 1-3 for both vertical and birdcage body coils. Table I (Figure 4) gives the maximum of 1g-averaged SAR around tips of left and right DBS leads when the input power of both coils was adjusted to generate the same flip angle 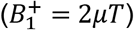 on a circular plane below the tip of the leads. For patient 1 with isolated leads, SAR amplification was reduced by 14-fold and 30-fold around tips of left and right DBS leads respectively, when exposed to RF fields of the vertical coil compared to the birdcage coil.

**Figure 4:**
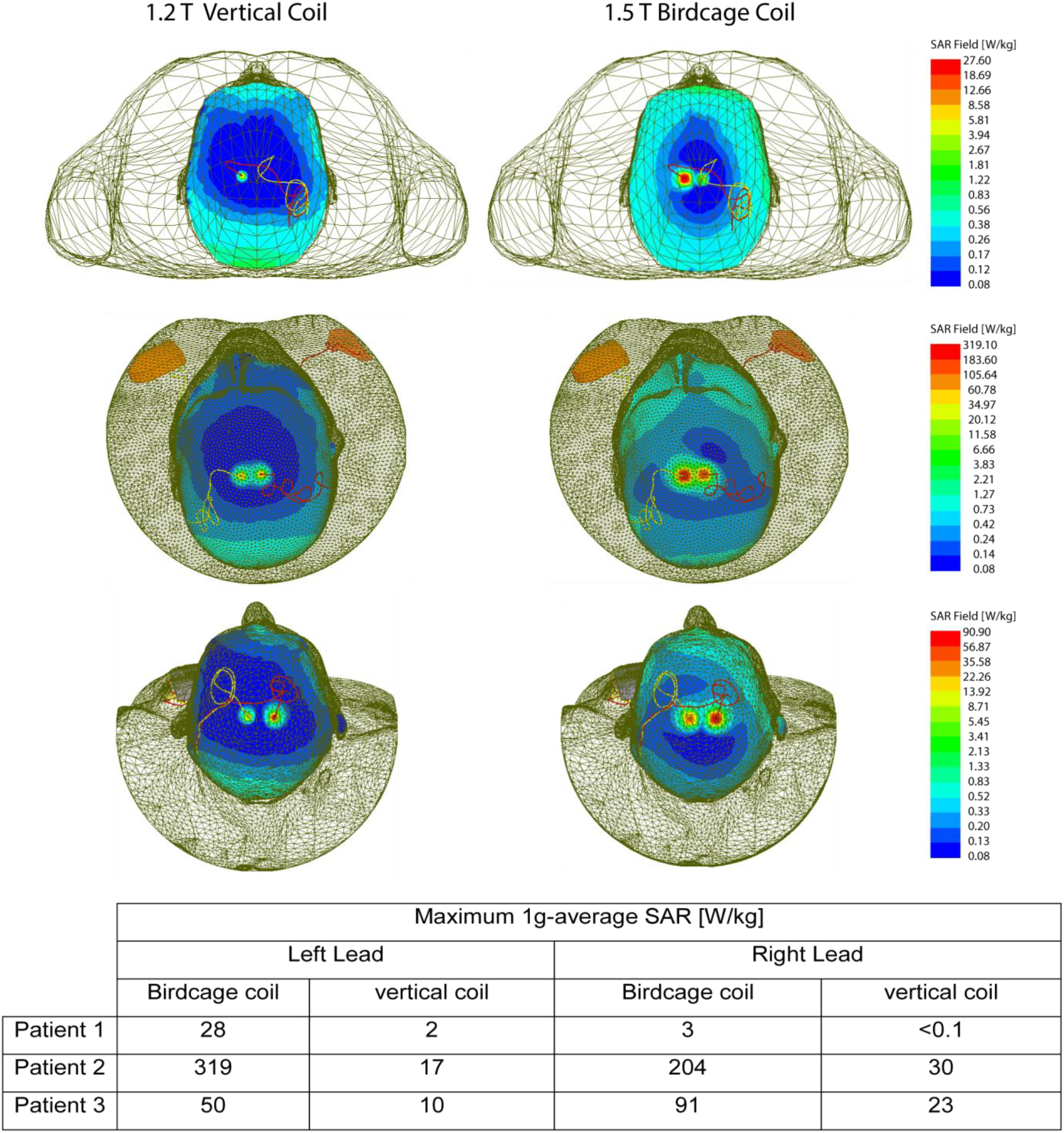
1gSAR distributions in patients 1-3 on an axial plane passing through distal electrode contacts for the vertical coil (1.2T) and birdcage coil (1.5T). The input power of both coil is adjusted to produce a mean B1+=2µT on a circular axial plane passing 20mm below the electrodes.

A substantially higher local SAR amplification was observed around tips of DBS leads in patient 2 who had a fully-implanted system with two bilateral IPGs. Compared to the birdcage coil, the vertical coil reduced the local SAR by 19-fold SAR at the tip of left DBS lead, and by 7-fold SAR reduction at tip of right lead.

Patient 3 represented a special case where concentric loops were introduced in the trajectory of the leads to reduce SAR [23, 27]. This technique works by cancelling out the effect of tangential component of the electric field along opposite sides of the loop [23]. Although due to this surgical modification patient 3 already demonstrated a reduced SAR at tips of left and right leads in the birdcage coil, vertical coil still further reduced the SAR by 5-folds at the tip of left lead and by 4-folds at the tip of right lead.

Figure 5 illustrates the orientation of the incident electric field vector along the trajectory of the leads in patient 1. As is can be observed, the electric field of vertical coil along DBS leads is oriented in anterior-posterior direction which is orthogonal to the trajectory of leads that roughly runs along medio-lateral direction on the scalp. In contrast, electric field of the birdcage coil has a significant tangential component along the extracranial portion of the leads, inducing a strong virtual voltage on lead wired which in turn generate RF currents. The video in the supplementary data shows the evolution of incident electric field along trajectories of the implants in time.

**Figure 5:**
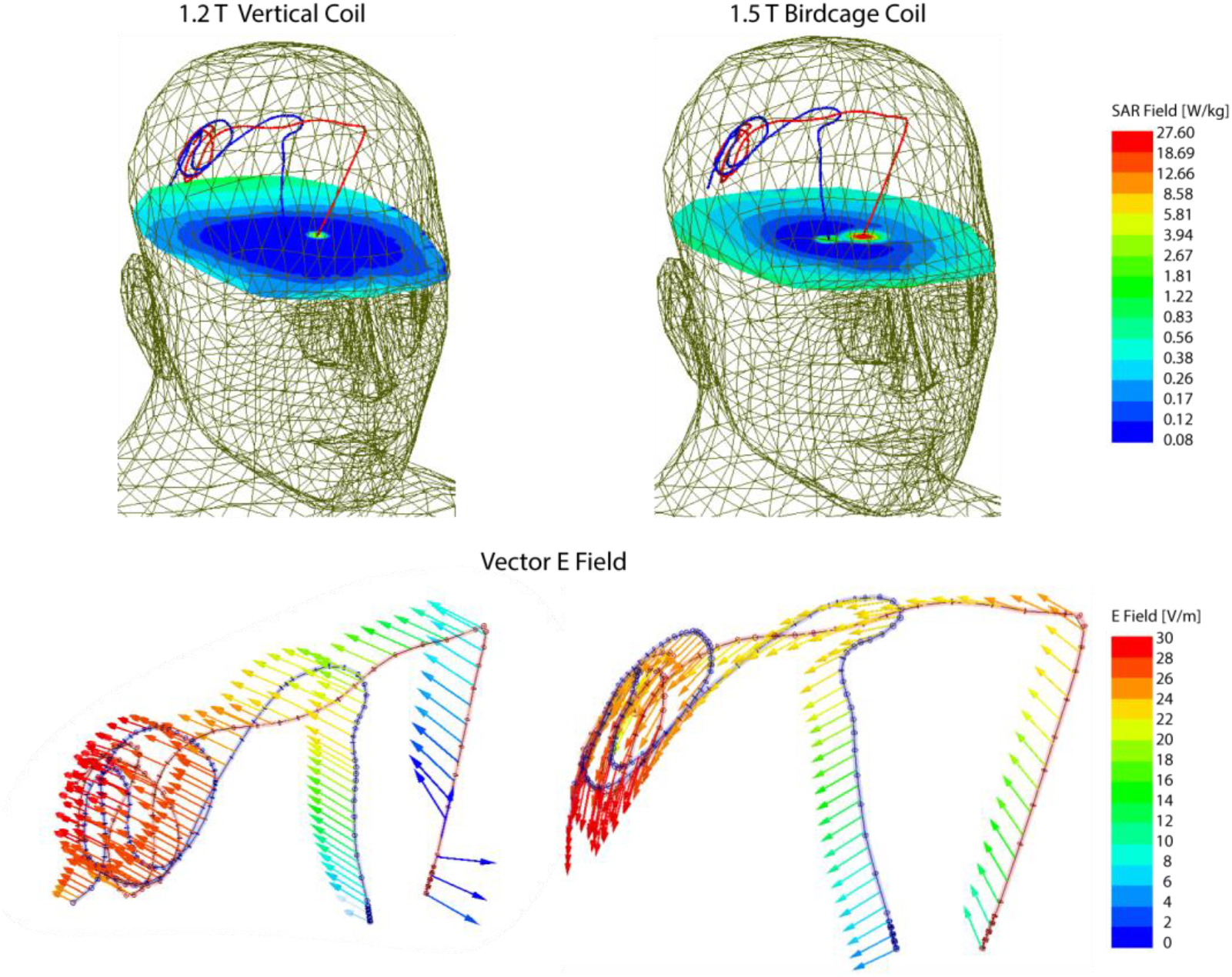
1gSAR and the vector E field in patient 1 for birdcage coil and vertical coil. The orientation of E vector is orthogonal to the extracranial trajectory of the lead in the vertical coil, minimizing the induction of RF currents. The time evolution of E vector along the trajectory of the leads is given in the Supplementary Data.

## Discussion and conclusion

During the past decade, DBS has developed into a remarkable treatment for major disabling neurological and psychiatric disorders [32–40], yet our knowledge of its underlying mechanisms of action remains rudimentary. The underlying affected brain circuits are largely unknown, undermining certainty in the optimal structures to target and most efficacious protocols. MRI of DBS patients is extremely useful to rule out complications, assess comorbidities, and interpret therapeutic effects of the stimulation. There are several situations when patients with actively implanted DBS devices need MRI. A majority require it as a part of diagnostic workup for various conditions such as acute ischemic stroke or postoperative hemorrhage [25, 41, 42]. The second scenario involves postoperative evaluation to document electrode location. This is particularly important considering the accuracy of implantation, defined as the difference between intended (or planned) and the actual (or implanted) electrode coordinates play an important role in determining the outcome [43, 44]. Current neuroimaging techniques for DBS electrode localization are mostly based on the co-registration of post-operative computed tomography (CT) and pre-operative MRI, with limited accuracy and cumbersome post-processing. Recent techniques for direct localization of individual DBS electrodes by MRI using zero-TE phase images demonstrated great spatial accuracy in phantom experiments [45] and their application in humans is highly desirable. Finally, for research applications, MRI is the modality of choice, especially for emerging neuromodulation indications such as mood and cognition disorders [46].

The main obstacle in application of MRI in patients with DBS implants is the RF heating due to coupling of scanner’s electric fields with implanted leads. Such concerns have led many centers to refrain from performing MRI on DBS patients mainly due to difficulty of complying with industry-proposed labeling of the device [47]. In some cases, patients have faced the proposition of explanting their neurostimulator to receive a diagnostic MRI [48]. Past few years have witnessed a spike in efforts to alleviate the problem of MRI-induced implant heating. Majority of work has been focused on modification of implant’s geometry, structure, or material [49–58]. Despite a spate of patents filed over the past two decades however, there is not a single MR-safe DBS product available in the market, attesting to the fact that the problem of RF safety cannot be addressed by device manufacturers alone. In response, several groups have worked on MRI hardware modification to reduce the antenna effect by shaping and steering electric fields of scanners. Techniques based on parallel transmit pulse tailoring [21, 28, 59–66] and reconfigurable MRI [18, 21, 67–71] have shown promising results in phantoms studies at 1.5T and 3T, but such techniques require sophisticated hardware setup and high level expertise to deploy and thus their application is likely to be limited to research rather than everyday clinical use. Vertical MRI systems were originally introduced as open-bore scanners offering an ideal platform for interventional procedures including direct DBS targeting with the help of interaoperative MRI [72–75]. The current MR labeling of DBS devices, as well as the entirety of MRI studies on the RF heating of conductive implants has been limited to horizontal (closed-bore) MRI systems. No DBS SAR literature exists for vertical MRI scanners which generate a fundamentally different electric and magnetic field distribution.

This work presents, for the first time, a computational study of RF heating around tips of DBS implants during MRI in a vertical open-bore system at 1.2T and compares the results with the SAR generated by a conventional 1.5T birdcage coil. Realistic models of DBS devices were constructed from postoperative CT images of patients, representing three major clinically-relevant device configurations. This included a patient with isolated bilateral leads, a patient with fully-implanted bilateral DBS leads stimulated via two single-channel implantable pulse generators (IPGs) implanted in right and left pectoral regions, and a patient with fully-implanted bilateral leads stimulated through one double-channel IPG implanted unilaterally. We found a significant reduction in SAR amplification (10-30 fold) at the tips of implanted leads in the case of isolated bilateral leads, and a 4-19-fold reduction in SAR in case of fully-implanted systems for vertical radial coil compared to the horizontal birdcage coil. A closer inspection of electromagnetic fields revealed that the orientation of incident electric field of vertical coil was mostly orthogonal to the trajectory of DBS leads, a criterion that we recently demonstrated to play a role in the severity of RF heating [23]. These results, if replicated in larger simulation cohorts and verified experimentally, can open the door to plethora of structural and functional MRI applications to interpret and guide DBS therapy. Interestingly, open bore MRI was originally developed to facilitate interventional applications, offering an ideal platform for MR-guided neurosurgery which can potentially revolutionize DBS surgery.

The problem of MRI-induced RF heating of medical implants is encountered in many other clinical applications. For example, integrated intracranial EEG (iEEG) with functional MRI is desired to elucidate mechanisms underlying the generation of seizures, yet temperature rises up to 10ºC has been reported in the tissue around iEEG electrode grids during MRI [76]. Many patients with MR-conditional cardiovascular electronic devices happen to have retained cardiac leads, which are contraindication for MRI [77, 78]. Our results also encourage future studies to assess the RF heating of other types of elongated implants in vertical MRI systems.

## Acknowledgement

This work has been supported by NIH grant R03EB025344, R03EB024705, and R00EB021320.

